# Metabolic modelling of the human gut microbiome in type 2 diabetes patients in response to metformin treatment

**DOI:** 10.1101/2022.04.22.489154

**Authors:** Bouchra Ezzamouri, Dorines Rosario, Gholamreza Bidkori, Sunjae Lee, Mathias Uhlen, Saeed Shoaie

## Abstract

The human gut microbiome has been associated with several metabolic disorders including type 2 diabetes. Understanding metabolic changes in the gut microbiome is important to elucidate the role of gut bacteria in regulating host metabolism. Here, we used available metagenomics data from a metformin study, together with genome-scale metabolic modelling of the key bacteria in individual and community-level to investigate the mechanistic role of the gut microbiome in response to metformin. Individual modelling predicted that species that are increased after metformin treatment have higher growth rates in comparison to species that are decreased after metformin treatment. Gut microbial enrichment analysis showed prior to metformin treatment pathways related to the hypoglycemic effect were enriched. Our observations highlight how the key bacterial species after metformin treatment have commensal and competing behavior, and how their cellular metabolism changes due to different nutritional environment. Integrating different diets showed there were specific microbial alterations between different diets. These results show the importance of the nutritional environment and how the dietary guidelines may improve drug efficiency through the gut microbiota.

## 1. Introduction

Type 2 diabetes mellites (T2DM) is a health burden with a rise in epidemic prevalence worldwide ^1^. T2D is characterized by increased blood glucose levels (hyperglycemia) ^2^. Metformin, is the most-prescribed medication to treat patients with T2DM due to its glucose lowering effects ^3^. Metformin propagates insulin sensitivity by mainly reducing hepatic glucose production (through an activation of the hepatic AMP-activated protein kinase protein) ^4^. The most common side effect of metformin is gastrointestinal discomfort including diarrhea, nausea, flatulence and bloating ^5^. There is growing evidence in animal and human studies suggesting that the gut microbiome is another target involved in the anti-diabetic effects of metformin ^6–9^. Recent investigations documented the therapeutic benefit of orally-administrated metformin compared to intravenously-administrated metformin in T2D patients, suggesting the beneficial contribution of the gut microbiota ^10^. Metformin alters the gut microbiome by enhancing *Escherichia sp, Akkermansia muciniphila* and *Subdoligranuum variable*, reducing *Intestinibacter bartletti* and increasing the levels the SCFAs; butyrate and propionate ^7,8^. This could indicate the anti-obesity property of metformin by modulating the gut microbiome and its metabolites, however the precise mechanisms are unclear.

Understanding the role of bacterial-derived gut metabolites can provide a platform to elucidate interactions between microbe-microbe, microbe-diet and drugs. The gut microbiota is an attractive target for therapeutic intervention and using nutrition may help to promote drug efficiency and reduce gastrointestinal side effects ^11^. To elucidate these interactions individual and systems level analysis is needed ^12–14^. Hence, systems biology approaches could be applied to reveal these associations between the abundances of different microbes and the molecular mechanisms underlying metformin treatment on a metabolic level ^15^. Genome-scale metabolic models (GEMS) have been used to gain a detailed understanding of microbial metabolic changes in various environments. Previous studies used GEMs to understand the metabolic interactions between microbes and host-microbes. ^16–18^.

Wu et al 2017 collected fecal samples from treatment naïve individuals that received 1,700 mg/d metformin treatment (22 patients) for 4 months and generated shotgun metagenomics data to determine the species abundances. In the present study we re-analyzed this metagenomics data with an updated gut microbial gene catalog and metagenome species profile ^19^. We analyzed the carbohydrate-active enzymes of the significantly altered species and found that species that are decreased after 4 months of metformin treatment showed an increased number of annotated mucins and host glycan degradation in comparison to the significantly increased species. We performed constraint-based analysis using GEMS integrating different diets to predict the phenotype of the drug metformin on the human gut microbiota. These diet-microbiota interactions can help us understand how to increase drug efficiency or mimic drug effects in the gut microbiome of patients with a dysbiosis to an improved phenotype.

## 2. Results

### 2.1 Gut microbiome profiling

Publicly available metagenomics data of 22 patient with treatment-naïve type 2 diabetes was quantified. In the metagenomics analysis, the latest available human gut microbiome catalog was used to determine the gene abundance. We used metagenome species pangenomes (MSPs) to identify gut microbial species that were perturbed with more in depth functional annotation of key metagenome species (Method). Here, the microbiome compositional changed between M0 (before metformin treatment) and M4 (after metformin treatment). The significant taxonomic changes on phylum, family and genus level between M0 and M4 can be found in supplementary table S1. Enterotype analysis was performed using a supervised clustering method with 3 clusters: Bacteroidetes, Firmicutes and Prevotella enterotypes (Figure 1A). We observed that before metformin treatment (M0) patients were clustered into Bacteroidetes and Firmicutes. After metformin treatment (M4) the microbiome changed to Firmicutes and Prevotella enterotype. Personalized analysis of the 22 patients showed how each patient enterotype changed after 4 months of metformin treatment (M4) (Figure 1B). The species-level abundances (Wilcoxon signed-rank test, false discovery rate (FDR <0.05) of 72 MSPs were significantly different between M4 and M0; 59 MSPs were significantly decreased and 13 significantly increased (Supplementary table 1). Using MSPs and the gut microbial catalog identified additional species with different abundances between M4 and M0 (Figure 1C). In accordance with the original study, the abundances of *Akkermansia muciniphila, Escherichia coli, Bifidobacterium* were increased whilst the abundance of *Intestinibacter bartlettii* was decreased. We identified additional species - *Blautia wexlerae* (a, short chain fatty acid producing species) with significantly higher abundances after metformin treatment and several species that had significantly decreased abundances such as *Alistipes obesi (Alistipes genera), Roseburia sp. CAG:100, Faecalibacterium prausnitzii 7, Faecalibacterium prausnitzii 3 (L2-6)*, butyrate producers, and several firmicutes bacteria. To better understand the interactions between the top 10 most significantly increased and decreased classified MSPs we constructed an integrative correlation network (ICN) ^31^. Before metformin treatment many negatively and positively correlated interactions exist between the different species. For instance, *Blautia wexleria* has a positively correlated interaction with the different *Clostridum* species whilst after metformin treatment this is lost. Moreover, after metformin treatment there are less interactions between species. (Figure 1D).

**Figure 1.**
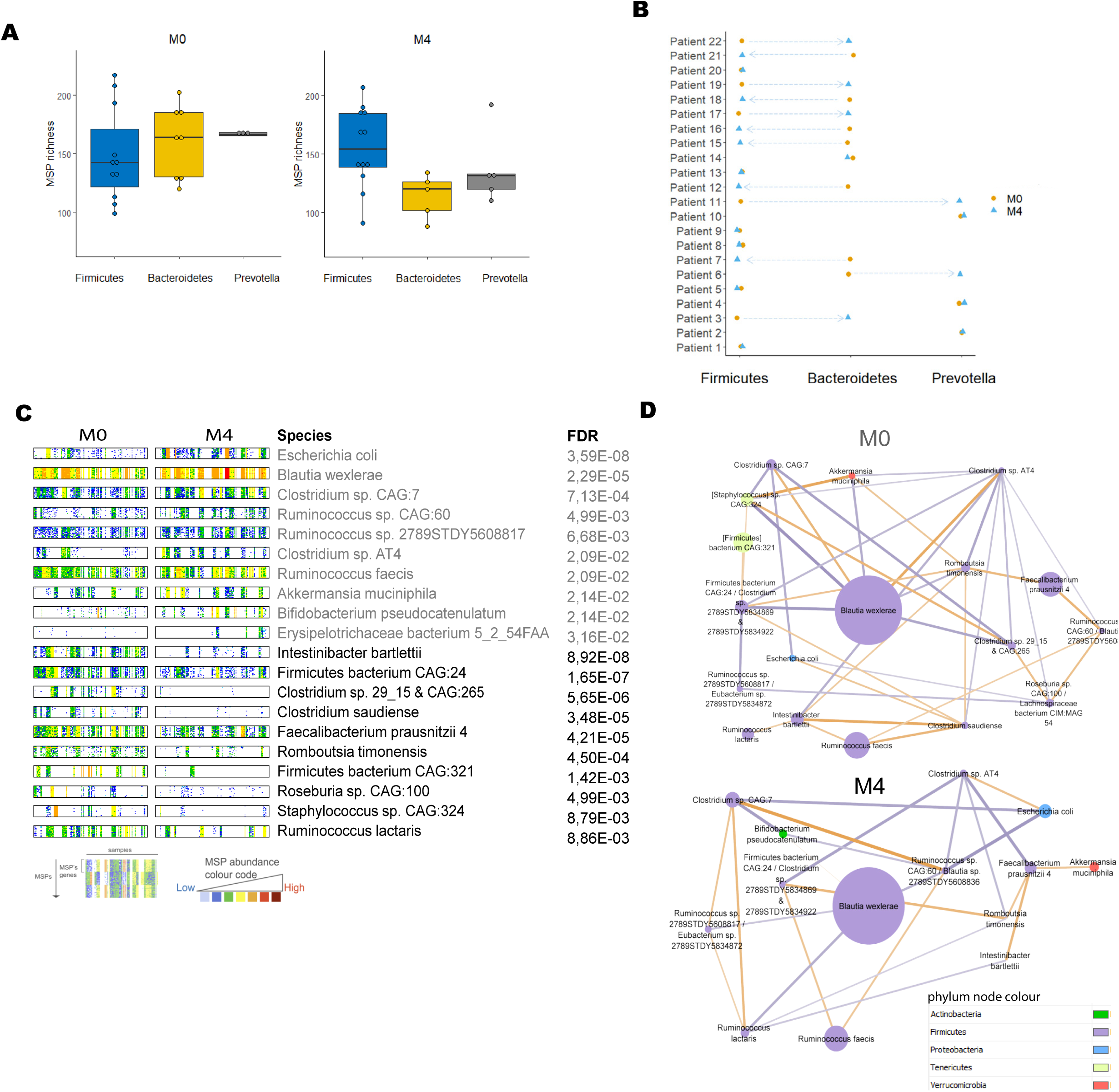
Enterotype and microbial compositional changes. (A) Relation between MSP richness and enterotype before and after metformin treatment (M0 and M4). (B)Personalized enterotype for each patient before and after metformin treatment (M0 and M4) (B) Barcodes of significantly increased and decreased (Wilcoxon signed-rank test, FDR < 0.05) MSPs between M4 and M0. (C) Barcodes represent each sample in the group and respective gene counts of each gene clustered into MSPs. Sample size under analysis: 10 samples that are significantly increased and decreased after metformin treatment (M4) (D) Integrative correlation network (ICN) based on MSPs significance and Spearmen correlation between M4 and M0 (Wilcoxon signed-rank test, FDR < 0.05). In the network, the orange edges correspond with a positive correlation whilst the purple edges correspond with a negative correlation. The nodes are coloured based on phylum level whilst the node size correspond with the abundance of each species.

### 2.2 Metabolism of carbohydrate was changed with metformin treatment

One of the key factors in the gut microbiome is the metabolism of carbohydrates. CAZymes analysis was performed on genes to identify potential differences in the microbial conversion of carbohydrates (Supplementary table 2). Overall, we found MSPs with increased abundance in pectins and mannose, whilst species that are decreased were associated with an increased number of multiple polysaccharides, mucins and host glycan degradation gene coding CAZymes (Figure 2A). The species *Akkermansia muciniphila* which is well known for degrading host glycans and mucans was significantly increased after metformin treatment.

**Figure 2.**
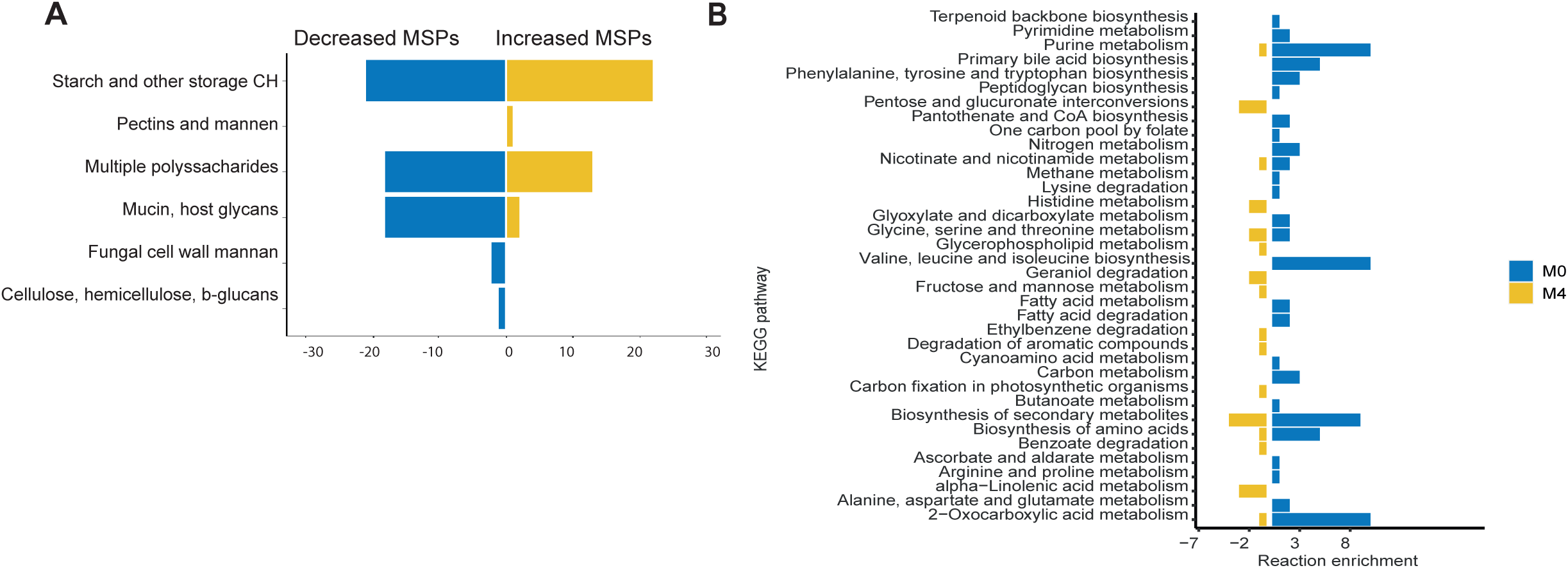
Functional annotation and reaction abundance pathway enrichment. (A) Carbohydrate-active enzymes (CAZymes) genes annotation of the top 10 most significantly increased and decreased species after metformin treatment (Wilcoxon signed-rank test, FDR<0.05). Genes annotated to mucin and host glycans are higher in species that are decreased after metformin treatment. (B) Significant metabolic pathway enrichment based on GEMs for MSPs in 22 patients before (M0) and after treatment (M4) (Wilcoxon signed-rank test, p<0.05).

### 2.3 GEMS for pathway analysis of associated MSPs

To understand the gut microbiota metabolism after metformin treatment and reveal interactions between the microbial species and the host, we downloaded the significantly increased and decreased GEM-MSPs with significantly altered abundances between M0 and M4 (https://www.microbiomeatlas.org/). These models were used for personalized community-level metabolic modelling and in depth pathway analysis. For each patient a matrix of reaction abundances before and after metformin treatment was generated based on MSP abundances and reactions that are present in the GEMs. The Kyoto Encyclopedia of Genes and Genomes (KEGG) metabolic pathways was used to identify the differences between the prevalence of metabolic reactions ^32^. Personalized metabolic pathway enrichment was performed using the reaction abundance matrix. Through KEGG pathway enrichment analysis, we identified significant alterations before and after metformin treatment (p < 0.05). Pathways such phenylalanine, tryptophane and tyrosine biosynthesis were enriched before metformin treatment. Moreover, we found that valine, leucine and isoleucine synthesis was enriched before metformin treatment. After metformin treatment we found enrichment in pathways such as pentose and glucuronate interconversions and histidine metabolism (Figure 2B).

### 2.4 GEMS of individual species associated with metformin treatment on various diets

Constrained-based analysis was used for the selected GEM-MSPs. To assess bacterial growth rate and major metabolic activities we used flux balance analysis (FBA) on each MSP-GEM model. Based on the FBA we assessed the potential contributions of bacteria to the gut metabolism. Each model was constrained based on their abundance and different diets (high fiber omnivorous (HFO), high fiber plant based (HFP), high protein omnivorous (HPO), high protein plant based (HPP), ketogenic (KETO) and western diet (WESTERN.) Overall, the simulations showed lower bacterial growth rate in all diets for species that were decreased in comparison to the increased species after metformin treatment (Figure 3A). We further analyzed the consumption and production of each metabolite for the increased and decreased species after metformin treatment. There was specific microbial alterations between each diets (Figure 3B, Supplementary figure 1-2). For instance, species that were increased produced a higher amount of butyrate on a HFO diet. Proline was produced by increased species after a HPP diet and KETO diet whilst arginine was produced on a HPO, HPP and HFP diet. Moreover, butyrate and propionate production was linked to decreased species on a HFP and HPO diet whilst increased species produced butyrate on a HFO and HPP diet. Additionally, the simulations showed that NH3 (ammonia) production is more present in species decreased after treatment on a HPP, HPO, Keto, and HFP diet. Increased species produce hydrogen sulphide (H2S) on all diets except the WESTERN diet.

**Figure 3.**
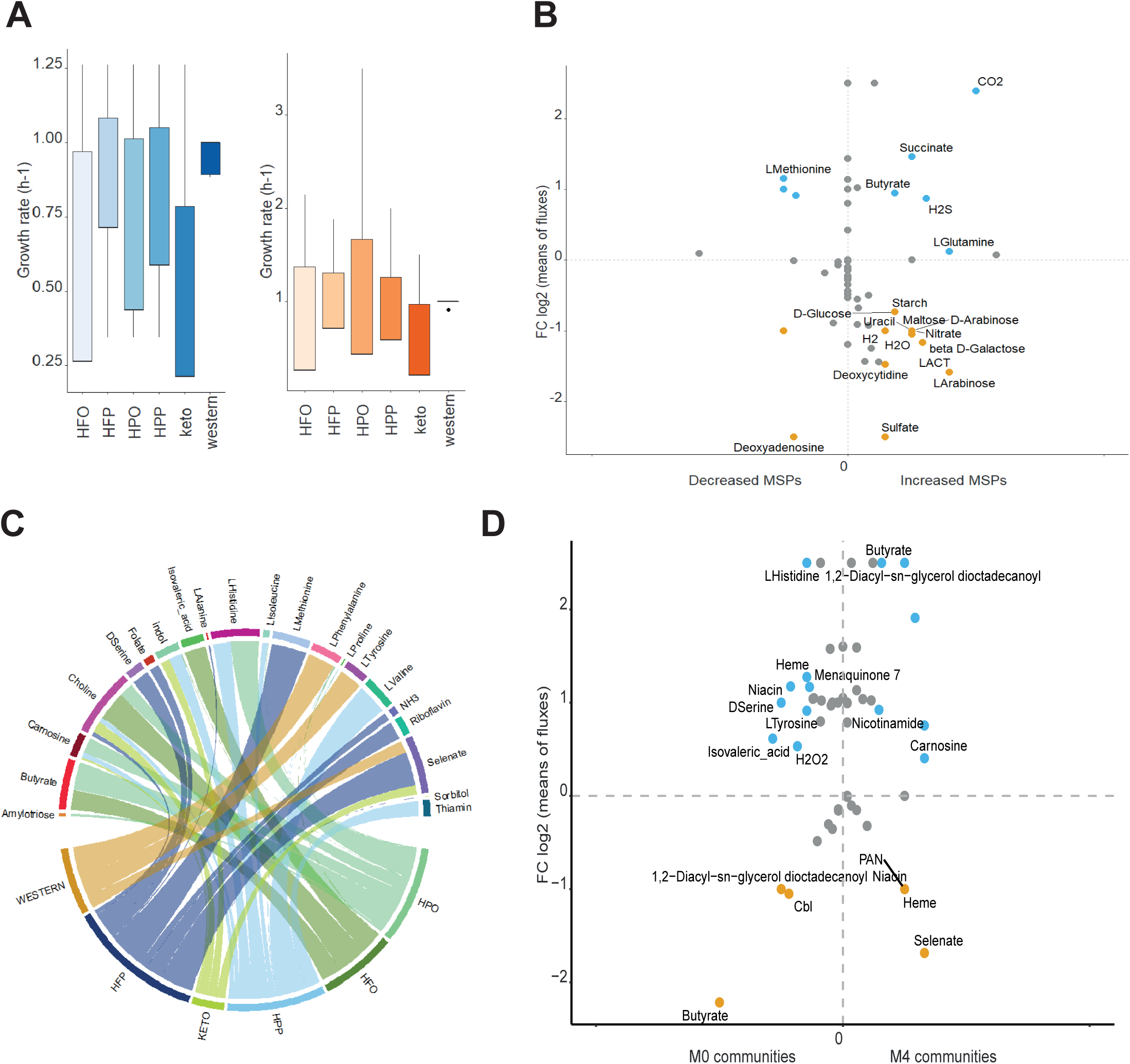
GEM of individual MSPs and personalized gut microbial community modeling. (A) Predictions of individual bacterial growth rate (h-1) constrained on different diets. Overall, MSPs with significantly decreased abundance (boxplots shown in blue) revealed a lower growth rate in comparison to MSPs with significantly increased abundance (boxplots shown in orange). (B) Potential contribution to host-intestinal metabolic pool based on metabolites production and consumption of significantly increased and decreased MSPs after metformin treatment (Wilcoxon signed-rank test, FDR < 0.05). The x axis represents the number of decreased or increased MSPs contributing to the metabolite consumption or production in y axis (orange negative or blue positive values, respectively). (C) Personalized gut microbial community modeling of each subject after metformin treatment (M4). Significantly (Wilcoxon signed-rank test, p < 0.05) different fluxes (FBA-based predictions) of secreted microbial metabolites on different diets are shown between gut-community models of patients after metformin treatment. The Chord diagram shows the significant means of microbial-metabolites fluxes on each diet. (D) Potential contribution of gut microbiota to host intestinal metabolic pool, based on personalized gut microbial community modeling. Increased secretion of microbial metabolites in controls communities compared to M4 communities are shown in blue; and orange represents the consumed microbial metabolites.

### 2.5 Metabolic community models show the gut microbial contribution to a changed metabolism after metformin treatment and the importance of diets

The individual microbial activity shows the potential role of each species in the overall gut microbiome metabolism. However, as the microbial species live in communities their behavior changes due to the availability of nutrients and the abundances of the species. Hence, we reconstructed personalized community models for each individual that obtained metformin treatment (22 patients in total) which allowed us to get a deeper insight in the contribution of the gut microbiota to host metabolism. For each personalized community a top 10 of most significantly increased and decreased MSPs per patient was used and different diets were constrained with community biomass as cellular objective. The significant secreted microbial metabolites on different diets before and after metformin treatment are shown in Figure 3C. The personalized community modelling constrained for a HFO diet showed that the metabolites butyrate, carnosine, and nicotinamide (vitamin B3) are produced after metformin treatment (Figure 3D). Proline gets produced when constrained on a ketogenic diet as well as valine, and carnosine. There was an increased microbial production of indole and histidine after metformin treatment on a constrained HPO and HPP diet. On a constrained western diet thiamin is secreted after metformin treatment (Supplementary figure 1).

## 3. Discussion

In this study, we performed downstream analysis of the gut microbial abundance alteration before and after metformin treatment using the latest human gut microbiome catalog. This allowed us to identify different gut microbial species and strains. Systems biology approaches have been used to understand the relationship between host and microbiota and to understand biological processes in more details ^8,14,16,33^. Here, we used GEMs for the key microbial species that are changed after metformin treatment and performed reaction abundance pathway analysis, individual and community modelling integrating different diets. These analysis gave us insight into the metabolic contribution of the gut microbiota and the effect of metformin treatment in naïve-type2 diabetes patients. We observed increased abundance in *Akkermansia muciniphila, Escherichia coli*, and *Bifidobacterium* whilst the abundance of *Intestinibacter bartlettii* was decreased, which is in agreement with the original study ^7,8^. We identified additional species such as *Blautia wexlerae* (a short chain fatty acid producing genus) with significantly higher abundances after metformin treatment and several species that had significantly decreased abundances like *Alistipes obesi* (Alistipes genera), *Roseburia sp. CAG:100, Faecalibacterium prausnitzii 7, Faecalibacterium* and *prausnitzii 3* (L2-6). These decreased species belong to the phyla Firmicutes which is contrary to previous studies showing that Firmicutes are major producers of butyrate and are found to be increased after metformin treatment. However, these observed species might be attributed to the differences in strain specific function. A study in rats demonstrated that metformin treatment changed the intestinal microbiota and showed an increase in the abundance of SCFA producing Blautia ^34^. This corresponds with the increased abundance found for the species *Blautia wexlerae*. Moreover, a cross-sectional study showed that the species *Blautia wexlerae* help reduce inflammation in obesity and insulin resistance ^35^. Roseburia has been found to be decreased in metformin treated patients ^36,37^. Also, Faeacalibacterim and Roseburia have been found to be negatively associated with T2D, so it is plausible that metformin treatment decreased these species even more ^38^. CAZymes analaysis showed that the top 10 most significantly decreased species have mucins and host glycan degradation. Mucins are critical for the maintenance of a homeostatic balanced between the gut microbiota and the host. During mucin degradation, monosacharides and amino acids are released, resulting in nutrients for other gut bacteria. Hence, we expected this phenotype to be present in the increased species after metformin treatment. A possible explanation for this is that the samples present from faeces are different than within the mucosa, which leads to an overrepresentation of bacteria that degrade mucins ^39^. A deeper investigation is needed to open new perspectives on the relationship of these species with the host and its effect due to drug treatment. Personalized reaction abundance analysis showed that before metformin treatment the branch chained amino (BCAA) e.g. leucine, valine and isoleucine were increased. Various studies using metabolomics showed that BCAA are elevated in patients with T2D ^40,41^. Elevated BCAA can activate the mechanistic target of rapamycin (mTOR) complex 1/ribosomal protein S6 Kinase pathway which leads to insulin resistance or to a build-up of metabolites that affect the pancreatic islet β-cells ^42,43^. After metformin treatment there were no pathways that were enriched in the BCAA suggesting that metformin reduces the circulating levels of BCAA. A study by Sriboonvorakul et al., 2021 showed that BCAA were significantly lower in T2D patients treated with metformin compared to healthy controls ^40,44^. A study in mice fed a high fat diet showed that BCAA levels decreased following metformin treatment ^45^. Another study, showed how reducing dietary BCAA might be able to restore metabolic health in obese mice consuming a high-fat, high sugar ‘western diet’ ^46^. Our modelling showed that isovaleric acid is significantly altered after metformin treatment with a high fiber omnivorous diet (HFO). Moreover, the individual modelling showed that species decreased after metformin treatment produce BCAA on HFP diet leading to a lower circulation of BCAA. The personalized community modelling showed that before treatment BCAA were produced but not after treatment. Reported side effects of metformin are bloating and intestinal discomfort. These side effects are enhanced by gasses produced by the gut microbiota. Our observations show that species that are increased after metformin treatment produce hydrogen sulphide (H_2_S). Hydrogen sulfide may have several beneficial effects for both host and gut microbiome such as anti-inflammatory properties and protects the gastrointestinal tract ^47^. We also showed that the aromatic amino acid biosynthesis (phenylalanine, tyrosine and tryptophan) was present before metformin treatment. A study by Alqudah et al., 2021 showed that the plasma concentrations of phenylalanine and tryptophane were increased in T2D ^48^. This is consistent with the personalized metabolic modelling where we saw a production of tryptophan in a HFP and HPP and western diet. Sugars such as glucose and fructose are one of the most analysed carbohydrates in metabolomics studies and have found to have a positive association with T2D ^49,50^. Here, we show that on different diets the species that are increased after treatment consume these sugars, showing that due to metformin the blood sugars levels get decreased so that the human body is more sensitive to insulin. SCFA produced by the gut can lead to beneficial effects on several tissues such as liver, muscle and adipose tissue; thereby improving insulin sensitivity ^51^. The ability to produce butyrate is enhanced with patients treated with metformin and can be increased by diet ^37 52^. Here, we show with personalized community modelling that butyrate is consumed before metformin treatment and in the individual models that species that are increased after metformin treatment produce butyrate. In a HFO diet butyrate and carnosine is produced after metformin treatment. Carnosine (β-alanyl-L-histidine) has been suggested to improve insulin sensitivity in type 2 diabetes patients, and a growing evidence of animal studies indicate a protective role of carnosine in diabetes.^53–55^. Overall, our modelling observations give us insight into how bacteria can have commensal and competing behaviours and how different diets leads to the production and consumption of extracellular metabolites such as SCFA, sugars, aromatic amino acids and BCAA. Understanding how nutritional environments can regulate metformin through the microbiota and elucidating the metabolic pathways may help inform personal dietary guidelines based on the gut microbiota to maximize the drug effect and reduce the gastrointestinal side effects.

## 4. Materials and Methods

### 4.1 Processing and downstream analysis of metformin metagenomics data

Publicly available shotgun metagenomics data on a randomized, placebo-controlled, double-blind type 2 diabetes (T2D) study which consisted of 40 individuals that were treated with either a placebo or metformin was downloaded from the sequence read archive under the accession: PRJNA361402 ^7^. The metagenomics cohort have been composed by Illumina NextSeq 500 paired end sequencing runs belonging to patients with metformin and placebo. Here, Meteor ^20^ was used to generate a gene abundance profiling table. Reads were mapped to the gut catalog ^19^ and gene count table for each samples was produced. The R Package MetaOMineR was used for the normalization of the gene counts table and further downstream analysis ^21^. As the metagenomics data was affected by variability in sequencing depth the data was downsized to 4 million reads to reduce this effect as it might bias the downstream analysis. The latest Metagenomic Species Pan-genomes (MSPs) profile was used to determine the metagenome species for each sample. The abundance of the species were calculated using marker genes in each MSP.

### 4.2 Richness and enterotype

The gene richness is highly sensitive to sequencing depth, therefore the mean of the 4 million downsized reads was considered as the richness. To study the samples’ bacterial diversity, the MSP richness was obtained based on the MSPs abundance table. DirichletMultinomial ^22^ for Clustering and Classification of Microbiome Data was used to identify, enterotypes of the samples by defining three components to model and by providing genera count data.

### 4.3 Carbohydrate-active enZymes annotation

Gut catalog genes ^19^ respective to MSPs were found to be significantly altered considering metformin dysbiosis (M4 against M0) and annotated to dbCAN2 ^23^. Based on literature review, we identified substrate conversion from annotated Carbohydrate-Active enZymes (CAZymes) ^24–28^.

### 4.4 KEGG metabolic pathway analysis based on reaction abundance

The MIcrobial and personalized GEM, REactobiome and community NEtwork modeling (MIGRENE) toolbox (https://github.com/sysbiomelab/MIGRENE) ^29^ was used for reaction abundance and pathway enrichment analysis. Microbial reaction abundance per sample of the metformin metagenomics was determined based on each MSP identified per sample that we could originate a functional metabolic model. A reaction pool was generated, and reactions were multiplied by MSP abundance in each sample. Personalized reaction abundance was calculated based on by summing each reaction frequency within each individual-microbiota. Reactions with significantly different abundances were identified by Wilcoxon signed-rank test (p < 0.05) for dysbiosis. Significant reactions were mapped to respective KOs, or Enzyme Commission number (EC number) when KO was not available, which in turn were mapped to KEGG metabolic pathways. The metabolic pathway enrichment was performed based on microbial reaction presence retrieved from the metabolic models. Hypergeometric test was applied for pathway enrichment. Significant (p < 0.05) enriched pathways were identified using Wilcoxon signed-rank test.

### 4.5 Individual modelling simulation

For individual modelling the COBRA toolbox was used, and the top 10 most significantly increased and decreased MSPs-GEMs were selected for individual metabolic modeling. Diet constrains were performed using flux balance analysis with biomass as the cellular objective function. The microbial growth rate; consumption and production by the bacteria were determined using Flux balance analysis (FBA). The simulation of functional GEMs was then constrained based on multiple diets (high protein omnivorous, high protein plant based, high fiber plant based, high fiber omnivorous, western diet and ketogenic diet) in anaerobic conditions

### 4.6 Personalized gut-microbial community metabolic models

Community models for each sample in the metformin cohort were reconstructed using the MIGRENE toolbox ^29^. A maximum of 10 MSPs per community was reconstructed due to computational power requirements and to assure model functionality. The S matrices were combined in order for each microbe to have their own cellular compartment; each microbial cell has a compartment which represents the intestinal lumen where metabolites from food ingestion is present. Another compartment is present for secreted microbial-metabolites and remaining food-derived metabolites that are not consumed by microbes. These microbial metabolites can be absorbed and reach blood circulation or excreted and present in human faeces. For each community, each individual bacterium biomass function was constrained on the respective abundance in each specific sample. Community biomass is composed of all the different microbes present in the community. The community biomass was defined as the objective function. For the community modelling the top 10 most abundant MSPs in each sample was selected per community. FBA was performed with biomass as the objective function and the predicted fluxes of exchange reactions repressing food-derived metabolites (FoEx) and microbial derived metabolites (FeEx) were obtained. The communities were constrained based on multiple diets (high protein omnivorous, high protein plant based, high fiber plant based, high fiber omnivorous, western diet and ketogenic diet) in anaerobic conditions. To identify principal microbial producers, reactions of interest were defined for the community models. Models that had similar number of productions across communities in both groups (M0 and M4), were ignored, in order to identify main contributors to increased or decreased secretions after metformin treatment (M4). Only microbial models with production in at least 3 communities in the group were considered.

### 4.7 Quantification and statistical analysis

Statistical analysis was performed using R software v 4.0.4. Significant taxonomic alterations between M0 (before metformin treatment) and M4 (after metformin treatment) were tested by Wilcoxon signed-rank test and corrected for multiple testing using Benjamini-Hochberg false discovery rate (FDR), q-values (significance FDR <0.05) ^30^, which can be found in Figure 1. In figure 2 the significant different abundances and enriched pathways were identified by Wilcoxon signed rank test (p <0.05) for metformin dysbiosis.

## Author Contributions

S.S. conceived and supervised the study. S.L. performed mapping of raw sequence to the gut catalog and produced the gene count tables. D.R and B.E performed the microbiome analysis. B.E performed the modeling. G.B. refined the individual and community GEMs. B.E wrote the manuscript. S.S., G.B., M.U. and D.R. provided critical comments and feedback. All authors have revised and contributed to the final version.

## Data Availability Statement

No new data was generated as part of this study.

## Acknowledgments

We thank the Swedish National Infrastructure for Computing at SNIC through Uppsala Multidisciplinary Center for Advanced Computational Science (UPPMAX) under Project SNIC 2020-5-222, SNIC 2019/3-226, SNIC 2020/6-153. The authors thank Ceri Proffitt for her technical support with the personalized community models.

## Supplementary Figure Legend

**Supplementary figure 1. GEM of individual MSPs and personalized gut microbial community modelling on a western, HFP and Keto diet**.

Left of panel A,B and C: Potential contribution to host-intestinal metabolic pool based on metabolites production and consumption of significantly increased and decreased MSPs after metformin treatment (Wilcoxon signed-rank test, FDR < 0.05). The x axis represents the number of decreased or increased MSPs contributing to the metabolite consumption or production in y axis (orange negative or blue positive values, respectively). Right of panel A, B and C: Potential contribution of gut microbiota to host intestinal metabolic pool, based on personalized gut microbial community modeling. Increased secretion of microbial metabolites in controls communities compared to M4 communities are shown in blue; and orange represents the consumed microbial metabolites. The heatmaps shows the Z scores of the means of microbial metabolites fluxes.

**Supplementary figure 2. GEM of individual MSPs and personalized gut microbial community modelling on a HFO and HPP diet**.

Left of panel A and B: Potential contribution to host-intestinal metabolic pool based on metabolites production and consumption of significantly increased and decreased MSPs after metformin treatment (Wilcoxon signed-rank test, FDR < 0.05). The x axis represents the number of decreased or increased MSPs contributing to the metabolite consumption or production in y axis (orange negative or blue positive values, respectively). Right of panel A and B: Potential contribution of gut microbiota to host intestinal metabolic pool, based on personalized gut microbial community modeling. Increased secretion of microbial metabolites in controls communities compared to M4 communities are shown in blue; and orange represents the consumed microbial metabolites. The heatmaps shows the Z scores of the means of microbial metabolites fluxes.

## Supplementary Table Legend

Table S1. Taxonomic alterations of gut microbiotas between M4 and M0 by applying Wilcoxon signed-rank test and by correcting the p values for multiple testing applying Benjamini-Hochberg FDR. Significance level FDR < 0.05.

Table S2. Substrate conversion of functional annotated CAZymes.

## References

1. Khan, M. A. B. et al. Epidemiology of Type 2 Diabetes – Global Burden of Disease and Forecasted Trends. Journal of Epidemiology and Global Health 10, 107 (2020).

2. Sharma, S. & Tripathi, P. Gut microbiome and type 2 diabetes: where we are and where to go? The Journal of Nutritional Biochemistry 63, 101–108 (2019).

3. Song, R. Mechanism of metformin: A tale of two sites. Diabetes Care vol. 39 187–189 (2016).

4. Buse, J. B. et al. The primary glucose-lowering effect of metformin resides in the gut, not the circulation: Results from short-term pharmacokinetic and 12-week dose-ranging studies. Diabetes Care 39, 198–205 (2016).

5. Kinaan, M., Ding, H. & Triggle, C. R. Metformin: An Old Drug for the Treatment of Diabetes but a New Drug for the Protection of the Endothelium. Medical principles and practice : international journal of the Kuwait University, Health Science Centre 24, 401–415 (2015).

6. Napolitano, A., Miller, S., Nicholls, A. W., Baker, D. & Van Horn, S. Novel gut-based pharmacology of metformin in patients with type 2 diabetes mellitus (PLoS ONE (2014) 9, 7 (e100778) DOI: 10.1371/journal.pone.0100778). PLoS ONE 9, (2014).

7. Wu, H. et al. Metformin alters the gut microbiome of individuals with treatment-naive type 2 diabetes, contributing to the therapeutic effects of the drug. Nature Medicine 23, 850–858 (2017).

8. Forslund, K. et al. Disentangling type 2 diabetes and metformin treatment signatures in the human gut microbiota. Nature 528, 262–266 (2015).

9. Shin, N. R. et al. An increase in the Akkermansia spp. population induced by metformin treatment improves glucose homeostasis in diet-induced obese mice. Gut 63, 727–735 (2014).

10. Horakova, O. et al. Metformin acutely lowers blood glucose levels by inhibition of intestinal glucose transport. 9, 1–11 (2019).

11. Valdes, A. M., Walter, J., Segal, E. & Spector, T. D. Role of the gut microbiota in nutrition and health. BMJ 361, 36–44 (2018).

12. Mardinoglu, A., Boren, J., Smith, U., Uhlen, M. & Nielsen, J. Systems biology in hepatology: approaches and applications. Nature reviews. Gastroenterology & hepatology 15, 365–377 (2018).

13. Ezzamouri, B., Shoaie, S. & Ledesma-Amaro, R. Synergies of Systems Biology and Synthetic Biology in Human Microbiome Studies. Frontiers in microbiology 12, (2021).

14. Rosario, D. et al. Systems Biology Approaches to Understand the Host–Microbiome Interactions in Neurodegenerative Diseases. Frontiers in Neuroscience 14, 716 (2020).

15. Gu, C., Kim, G. B., Kim, W. J., Kim, H. U. & Lee, S. Y. Current status and applications of genome-scale metabolic models. Genome Biology vol. 20 1–18 (BioMed Central Ltd., 2019).

16. Shoaie, S. et al. Quantifying Diet-Induced Metabolic Changes of the Human Gut Microbiome. Cell Metabolism 22, 320–331 (2015).

17. Rosario, D. et al. Understanding the Representative Gut Microbiota Dysbiosis in Metformin-Treated Type 2 Diabetes Patients Using Genome-Scale Metabolic Modeling. Frontiers in Physiology 9, 775 (2018).

18. Tramontano, M. et al. Nutritional preferences of human gut bacteria reveal their metabolic idiosyncrasies. Nature Microbiology 2018 3:4 3, 514–522 (2018).

19. Wen, C. et al. Quantitative metagenomics reveals unique gut microbiome biomarkers in ankylosing spondylitis. Genome Biology 18, 142 (2017).

20. Pons, N., Batto, J., Kennedy, S., … M. A.-J. O. en & 2010, undefined. METEOR, a platform for quantitative metagenomic profiling of complex ecosystems. researchgate.net.

21. Le Chatelier, E. et al. Richness of human gut microbiome correlates with metabolic markers. Nature 500, 541–546 (2013).

22. Holmes, I., Harris, K. & Quince, C. Dirichlet Multinomial Mixtures: Generative Models for Microbial Metagenomics. PLOS ONE 7, e30126 (2012).

23. Zhang, H. et al. dbCAN2: a meta server for automated carbohydrate-active enzyme annotation. Nucleic acids research 46, W95–W101 (2018).

24. Baroncelli, R. et al. Gene family expansions and contractions are associated with host range in plant pathogens of the genus Colletotrichum. BMC Genomics 17, (2016).

25. Borin, G. P. et al. Comparative Secretome Analysis of Trichoderma reesei and Aspergillus niger during Growth on Sugarcane Biomass. PLOS ONE 10, e0129275 (2015).

26. Breier, M. et al. Targeted Metabolomics Identifies Reliable and Stable Metabolites in Human Serum and Plasma Samples. PLOS ONE 9, e89728 (2014).

27. Geisler-Lee, J. et al. Poplar Carbohydrate-Active Enzymes. Gene Identification and Expression Analyses. Plant Physiology 140, 946–962 (2006).

28. Wegmann, U. et al. Complete genome of a new Firmicutes species belonging to the dominant human colonic microbiota (‘Ruminococcus bicirculans’) reveals two chromosomes and a selective capacity to utilize plant glucans. Environmental Microbiology 16, 2879–2890 (2014).

29. Bidkhori, G. et al. The Reactobiome Unravels a New Paradigm in Human Gut Microbiome Metabolism. bioRxiv 2021.02.01.428114 (2021) doi:10.1101/2021.02.01.428114.

30. Benjamini, Y. Discovering the false discovery rate. Journal of the Royal Statistical Society: Series B (Statistical Methodology) 72, 405–416 (2010).

31. Rosario, D. et al. Systematic analysis of gut microbiome reveals the role of bacterial folate and homocysteine metabolism in Parkinson’s disease. Cell Reports 34, (2021).

32. Kanehisa, M. Toward understanding the origin and evolution of cellular organisms. Protein science : a publication of the Protein Society 28, 1947–1951 (2019).

33. Mardinoglu, A. & Nielsen, J. Systems medicine and metabolic modelling. in Journal of Internal Medicine vol. 271 142–154 (2012).

34. Zhang, X. et al. Modulation of gut microbiota by berberine and metformin during the treatment of high-fat diet-induced obesity in rats. Scientific reports 5, (2015).

35. Benítez-Páez, A. et al. Depletion of Blautia Species in the Microbiota of Obese Children Relates to Intestinal Inflammation and Metabolic Phenotype Worsening. mSystems 5, (2020).

36. Hiel, S. et al. Link between gut microbiota and health outcomes in inulin -treated obese patients: Lessons from the Food4Gut multicenter randomized placebo-controlled trial. Clinical Nutrition 39, 3618–3628 (2020).

37. Mueller, N. T. et al. Metformin Affects Gut Microbiome Composition and Function and Circulating Short-Chain Fatty Acids: A Randomized Trial. Diabetes Care 44, (2021).

38. Gurung, M. et al. Role of gut microbiota in type 2 diabetes pathophysiology. EBioMedicine 51, 102590 (2020).

39. Ouwerkerk, J. P., De Vos, W. M. & Belzer, C. Glycobiome: Bacteria and mucus at the epithelial interface. Best Practice & Research Clinical Gastroenterology 27, 25–38 (2013).

40. Xu, F. et al. Metabolic signature shift in type 2 diabetes mellitus revealed by mass spectrometry-based metabolomics. The Journal of clinical endocrinology and metabolism 98, (2013).

41. Wang, T. J. et al. Metabolite profiles and the risk of developing diabetes. Nature medicine 17, 448–453 (2011).

42. Chen, X. & Yang, W. Branched-chain amino acids and the association with type 2 diabetes. Journal of Diabetes Investigation 6, 369–370 (2015).

43. Olson, K. et al. Alloisoleucine differentiates the branched‐chain aminoacidemia of Zucker and dietary obese rats. Wiley Online Library 22, 1212–1215 (2014).

44. Sriboonvorakul, N. et al. Low branched chain amino acids and tyrosine in thai patients with type 2 diabetes mellitus treated with metformin and metformin-sulfonylurea combination therapies. Journal of Clinical Medicine 10, 5424 (2021).

45. Riera-Borrull, M. et al. Metformin potentiates the benefits of dietary restraint: A metabolomic study. International Journal of Molecular Sciences 18, (2017).

46. Cummings, N. E. et al. Restoration of metabolic health by decreased consumption of branched-chain amino acids. The Journal of Physiology 596, 623–645 (2018).

47. Wallace, J. L., Motta, J.-P. & Buret, A. G. Hydrogen sulfide: an agent of stability at the microbiome-mucosa interface. American Journal of Physiology-Gastrointestinal and Liver Physiology 314, G143–G149 (2018).

48. Alqudah, A., Wedyan, M., Qnais, E., Jawarneh, H. & McClements, L. Plasma Amino Acids Metabolomics’ Important in Glucose Management in Type 2 Diabetes. Frontiers in Pharmacology 12, 1786 (2021).

49. Yang, S. J., Kwak, S. Y., Jo, G., Song, T. J. & Shin, M. J. Serum metabolite profile associated with incident type 2 diabetes in Koreans: findings from the Korean Genome and Epidemiology Study. Scientific Reports 2018 8:1 8, 1–10 (2018).

50. Rebholz, C. M. et al. Serum metabolomic profile of incident diabetes. Diabetologia 61, 1046–1054 (2018).

51. van der Hee, B. & Wells, J. M. Microbial Regulation of Host Physiology by Short-chain Fatty Acids. Trends in Microbiology 29, 700–712 (2021).

52. Knudsen, K. E. B. et al. Impact of Diet-Modulated Butyrate Production on Intestinal Barrier Function and Inflammation. Nutrients 10, (2018).

53. De Courten, B. et al. Effects of carnosine supplementation on glucose metabolism: Pilot clinical trial. Obesity 24, 1027–1034 (2016).

54. Köppel, H. et al. L-carnosine inhibits high-glucose-mediated matrix accumulation in human mesangial cells by interfering with TGF-β production and signalling. Nephrology, dialysis, transplantation : official publication of the European Dialysis and Transplant Association - European Renal Association 26, 3852–3858 (2011).

55. Sauerhöfer, S. et al. L-carnosine, a substrate of carnosinase-1, influences glucose metabolism. Diabetes 56, 2425–2432 (2007).

