## Supplementary Figure 1 for "Metabolic modelling of the human gut microbiome in type 2 diabetes patients in response to metformin treatment"

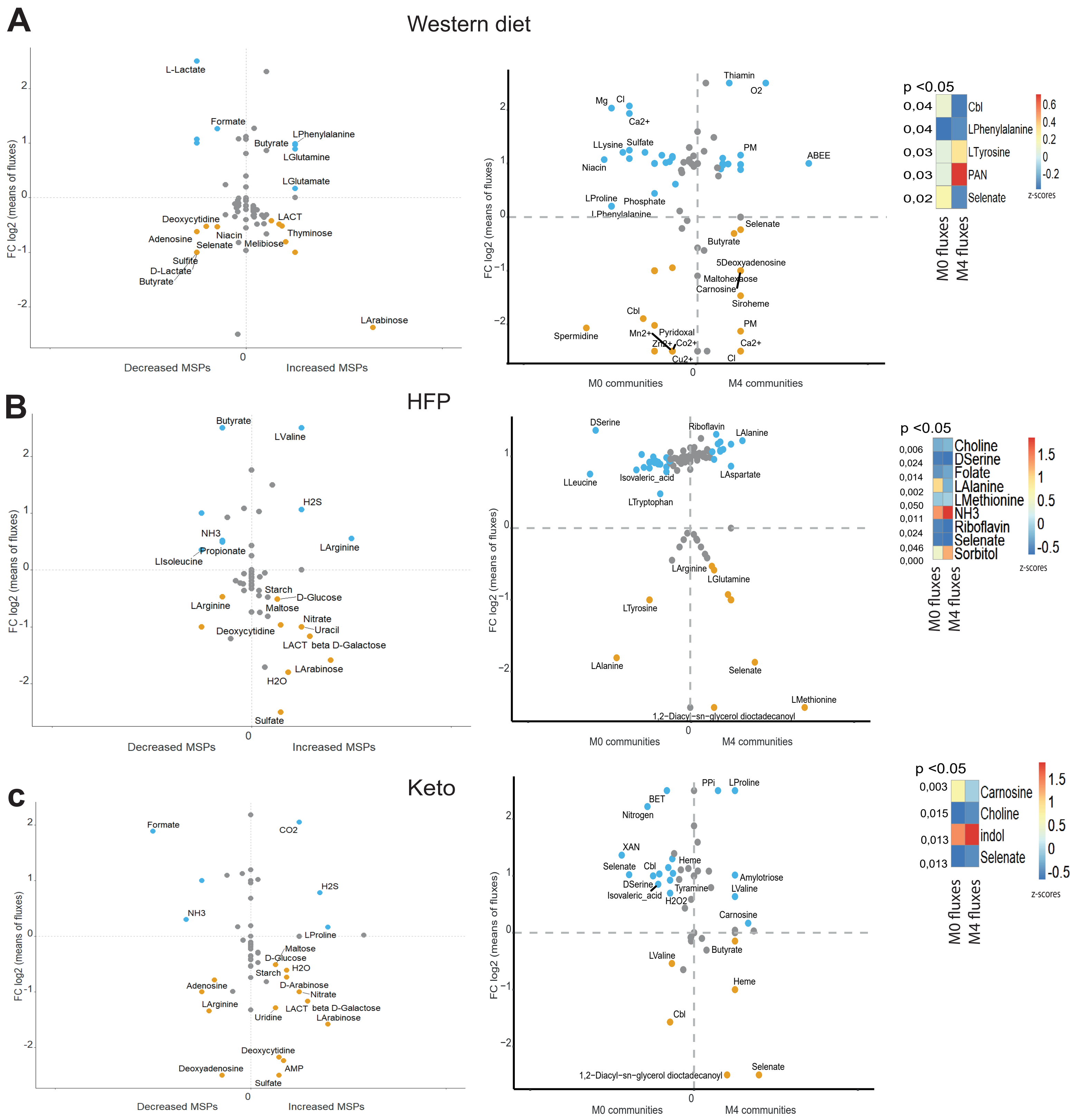


**Supplementary figure 1. GEM of individual MSPs and personalized gut microbial community modelling on a western, HFP and Keto diet.**

Left of panel A,B and C: Potential contribution to host-intestinal metabolic pool based on metabolites production and consumption of significantly increased and decreased MSPs after metformin treatment (Wilcoxon signed-rank test, FDR < 0.05). The x axis represents the number of decreased or increased MSPs contributing to the metabolite consumption or production in y axis (orange negative or blue positive values, respectively). Right of panel A, B and C: Potential contribution of gut microbiota to host intestinal metabolic pool, based on personalized gut microbial community modeling. Increased secretion of microbial metabolites in controls communities compared to M4 communities are shown in blue; and orange represents the consumed microbial metabolites.
